# Conditional source attribution at plankton bloom onset: Identifiability and sharp bounds under environmental forcing

**DOI:** 10.64898/2026.08.13.744670

**Authors:** Luigi Caputi

## Abstract

Can observations distinguish a bloom supplied from within a study volume from one supplied across its boundary? We develop a theoretical framework for that question at plankton bloom onset, conditional on a predeclared, observed or calibrated onset event and a declared set of environmental paths, biological responses, and model forms. The estimand follows source-event labels through forcing-dependent survival and genotype-specific growth. Its central certificate asks whether the local onset fraction is invariant over every source history that produces the same time-expanded observation record. For polyhedral history fibers, a Charnes–Cooper transformation computes both sharp dynamic-data endpoints as linear programs. When each source instead has a fixed normalized onset signature, the certificate reduces to a row-space test; uncertain signatures require a joint lifted program. For a finite compatible scenario ensemble, admissible fractions are the union across scenarios, and a point is justified only when every nonempty scenario gives the same singleton. A synthetic two-genotype witness gives the same observed total but local fractions of 2*/*3 and 1*/*3 under reversed forcing–response gains. The observer, mixture, and optimization ingredients are established; the contribution is their target-specific synthesis around source at onset. The framework is diagnostic rather than predictive. It specifies what a study must measure—local sources, boundary inflow, forcing, response, timing, and carrier signatures on one declared window—and returns an interval when missing components have justified bounds, including [0, 1] when they remain unconstrained.

## 1 Introduction

A bloom needs both a permissive environment and a population able to exploit it. Temperature, nutrients, and light can interact in their effects on phytoplankton growth [30]. Factorial shipboard experiments also show linked responses to temperature and nutrient supply [21]. These relationships are not universal constants: growth responses vary among and within taxa [5, 41].

Source availability answers a different question. Cells may persist inside a study volume, emerge from an internal reservoir, or cross an external boundary. Currents can deliver nonlocal populations [70, 71]. A pelagic rare biosphere can persist between favorable periods [25], while benthic resting stages provide a distinct reservoir mechanism [1, 2]. Neither observation explains a bloom by itself.

A fixed station can conflate these processes. Environmental measurements may show that growth was possible without showing where the responding cells entered. A repeated taxon name may show recurrence without preserving carrier identity. Conversely, a source panel may show that cells were available without showing that the environment allowed population expansion.

We therefore condition source attribution on a declared environmental path, biological-response specification, model form, and bloom-onset event observed through a predeclared channel. The target is the fraction of carrier-equivalent abundance at onset attributable to local admission events. Local events comprise the initial in-volume state and release from compartments inside the boundary. Imported events are inward crossings of the external boundary during the source window. Environmental conditions and carrier response after admission can change the fraction through differential survival or growth.

This target is most useful when bloom composition changes among events. The same taxon can contain populations with distinct genomic structure [26, 49, 62]. Closely related carriers can also have different thermal responses [41]. A change in genotype frequency may reflect a changed source mixture, post-entry selection, or both. The model below separates those explanations when the measurements permit it.

### 1.1 Contributions and scope

Functional observability, unknown-input geometry, row-space estimability, linear-fractional programming, and mixture perspective lifts are established mathematical tools. The contribution is their target-specific interdisciplinary synthesis: a source-at-onset estimand, its feasible dynamic history, explicit environmental–response scenarios, sharp partial-identification bounds, and a prospective measurement diagnostic.

Within that scope, the paper makes four contributions.

1. It defines a source-event onset fraction conditional on an explicit, system-specific scenario and an onset event tied to an observed or calibrated channel.
2. It uses classical functional-observability and unknown-input criteria to give a finite-horizon feasible-fiber certificate for that affine-ratio target. The shared-input-channel failure is a corollary.
3. It separates environmental-path, biological-response, and model-form uncertainty, gives exact linear programs for each dynamic-data endpoint, and gives sharp endpoint bounds and a width decomposition for finite ensembles.
4. It makes the full dynamic-history model primary, treats fixed onset signatures as a reduced special case, handles unknown provenance, and distinguishes a full-panel point-claim gate from bounded partial-identification usability.

The scope is conditional source-at-onset attribution rather than a general theory of bloom recurrence or genotype dominance. Environmental fit alone is not an origin certificate, and short-marker recurrence is not lineage continuity. Field point and interval claims follow the measurement and partial-identification gates stated below.

## 2 Scientific objects and conditional estimand

### 2.1 Five distinct questions

Table 1 separates observations that are often discussed as if they were interchangeable.

**Table 1.** Scientific objects kept separate in this paper.

| Object | Meaning |
| --- | --- |
| Taxon recurrence | The same named taxon is detected in more than one event. |
| Genotype recurrence | A declared multilocus or genome-wide carrier signature appears in more than one event. |
| Source-event contribution | Carrier-equivalent abundance at onset assigned to an initial state or admission event. |
| Post-entry selection | Forcing-dependent survival or reproduction changes the relative contribution after admission. |
| Genealogical ancestry | The longer-term lineage history of the carriers. It is not the target here. |

### 2.2 Boundary, source window, and carrier

Fix a three-dimensional control volume *B* and a source window [*t*_0_, *t*_⋆_]. The window ends at a bloom onset defined before source analysis. Vertical movement inside *B* is local. Entry across a declared external face is import. Re-entry after exit is a new imported event. Changing the boundary changes the estimand.

#### Definition 1

(Carrier and onset contribution). A *carrier* is the modeled unit propagated by the state equation. It may be a living cell, a genotype-resolved cell equivalent, or a calibrated biomass unit. A sequencing read is an observation, not a carrier, unless a validated conversion gives it a common carrier scale. An *onset contribution* is the amount of the carrier unit present at *t*_⋆_ and assigned to one source event under the declared propagation model.

#### Meaning

The target concerns contributions present at onset. It is not the unweighted number of cells that crossed a boundary earlier in the window.

Index modeled source–carrier classes by *j* = 1, …, *J*. Each class pairs one source-event label with one biological response class, such as a genotype. A physical source can contribute several values of *j*, and one genotype can occur under several source labels. Partition the classes into local L and imported I sets. This partition exhausts the modeled carrier mass. When geographic status is unresolved, the model uses paired latent local and imported components with a shared signature, as described below. It never assigns that mass to one side for convenience. The labels are window-specific source-event labels, not mutually exclusive genotype names: a genotype present initially and the same genotype imported later occupy distinct classes even when they share biological response parameters and an assay signature.

### 2.3 Forcing and bloom onset

Let *F*_t_ be the environmental and ecological forcing state. Its components must be chosen for the biological system. Possible measured components include temperature, dissolved nutrients, irradiance, salinity, and water-column mixing [13, 21, 30, 58]. Biological controls may be included when primary evidence makes them material. No universal list is assumed.

The forcing record can contain three kinds of entries. Some components are observed directly. Some are estimated with a stated error model. Others are bounded or latent. We write

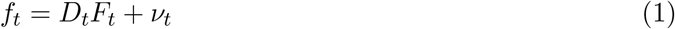

for environmental observations. Environmental-path uncertainty is

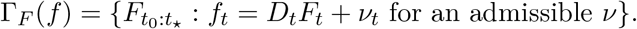

It is distinct from uncertainty in the response parameters or functions *β* = (*β*_1_, …, *β*_J_) and from model-form uncertainty. Let Γ_β_(*F*) contain response specifications supported by calibration or primary biology under path *F*, and let Γ ℳ (*F, β*) contain declared operator or model-form choices. The complete scenario set is

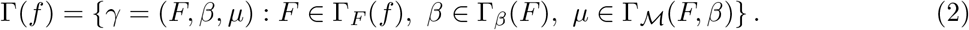

A known category is represented by a singleton. Thus uncertainty in a measured temperature path is not relabeled as uncertainty in genotype response, and uncertainty in a response curve is not hidden in an environmental error bar. The model-form index *µ* selects the declared propagation, entry, and observation operators and is suppressed in their notation below.

Let 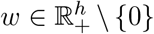 select abundance on the declared carrier scale, and let *N*_t_ = *w*^⊤^*x*_t_ be latent carrier abundance. An operational field event must be tied to an observed or calibrated onset channel,

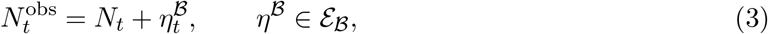

where ℰ_ℬ_ is a declared joint error set. The exact-calibration case has *η*^B^ = 0. One possible onset rule uses predeclared lower and upper thresholds, with 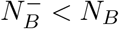:

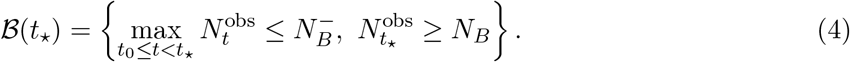

An application may use another operational rule. The rule, sampling cadence, units, observation operator, and error set must be fixed before source analysis. The onset-channel constraints enter the same feasible history set as the other observations. Equation (4) operationally locates onset rather than asserting its biological cause. A latent-state onset rule is model dependent and therefore belongs in *µ*. The closed thresholds also allow compact feasible fibers when the remaining state, error, and input bounds are closed and bounded.

A complete scenario is *bloom permitting* when at least one admissible state and source history reaches the predeclared event. The source target is evaluated over histories that reach that event under the declared initial-state and source-input bounds.

## 3 Forcing-dependent source model

### 3.1 Source-labeled population balance

Let 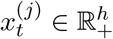 carry the composite source-event and biological-response label *j*, and let 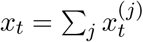 Membership in L or I records source geography; *β*_j_ records the genotype or other response class. For a fixed scenario *γ*, the labeled balance is

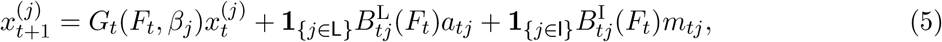

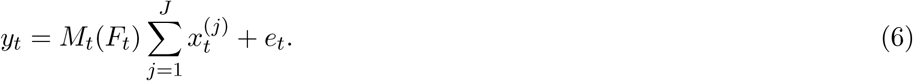

All source inputs are nonnegative. Because source geography is defined relative to *B* and [*t*_0_, *t*_⋆_], every carrier already inside *B* at *t*_0_ is initial local stock. We therefore impose the standing convention

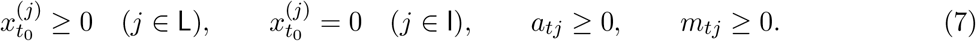

Thus any class with positive initial state belongs to L. If one genotype is present at *t*_0_ and is also imported later in the window, it is represented by two source-event classes, *j*_L_ and *j*_I_, which may satisfy 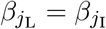 and share an assay signature but retain different geography and admission labels. This window-relative convention makes no claim that the ultimate ancestry of the initial carriers is local. Each transition and entry operator maps the nonnegative cone into itself, so a labeled carrier contribution cannot become negative.

The operator *G*_t_(*F*_t_, *β*_j_) represents survival, growth, activation-state change, loss, and within-volume redistribution. It may differ among genotypes. The matrices 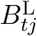 and 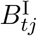 place local release and boundary inflow into the state. Forcing can affect entry survival or the state into which a carrier enters.

The model separates four roles. Environmental forcing affects whether growth is possible. Source inputs determine which carriers are available. The response parameters determine how each carrier class changes after entry. The model-form index selects the maintained operator family. This separation prevents environmental measurement error, biological response uncertainty, and structural model uncertainty from being reported as one undifferentiated “forcing” term.

### 3.2 When source labels can be propagated

Equation (5) uses additive source labels. It requires the following conditions.

1. A common carrier scale for all labeled contributions.
2. Nonnegative states, inputs, and positive-cone-preserving propagation.
3. Linear superposition of labeled states over the source window.
4. A fixed or explicitly varied complete scenario during each calculation.
5. Source-specific selection represented by *G*_t_(*F*_t_, *β*_j_) without density-dependent interaction among labeled states.
6. Stable labels, or a predeclared fractional inheritance rule, through mutation, life-history change, and reproduction.

Genotype-specific selection does not by itself violate additivity. It can be represented by different transition operators while labels remain traceable. Ecological interactions and priority effects can invalidate a source-mixture map [69]. Density-dependent transitions likewise break linear superposition. Cross-source mating or recombination instead breaks label stability unless a fractional inheritance rule is declared. In any of these cases, the linear recovery results do not apply without a justified extension. A study must use a nonlinear attribution rule or retreat to a source-flux target.

Forcing also need not be exogenous. If carrier abundance feeds back on nutrients, light, or grazer state during the source window, *F*_t_ must join the state equation. Treating the observed forcing path as fixed would then be an invalid simplification.

### 3.3 Conditional onset fraction

For *s > t*, define the source-specific propagator

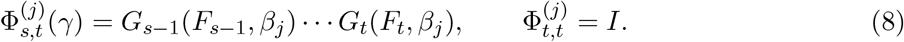

The local and imported onset contributions are

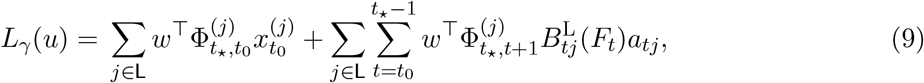

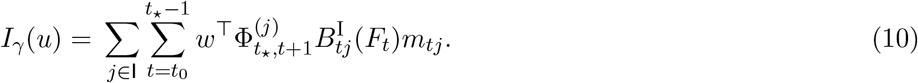

The vector *u* stacks the unknown source histories and any uncertain components of the local initial state. Known history components are kept fixed and enter the affine offsets below. The conditional target is

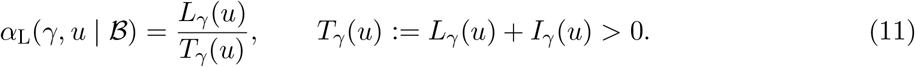

The positivity conditions give 0 ≤*α*_L_ ≤1. The event ℬ restricts the feasible histories. The complete scenario *γ* changes the weights assigned to entry histories through survival and growth. Thus equal entry fractions need not give equal onset fractions.

For source–carrier class *j*, define its unnormalized onset contribution and normalized share by

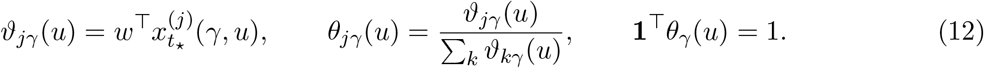

Expanding Equation (5) to *t*_⋆_ and using 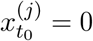 for *j* ∈ I gives 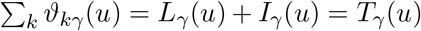.

With *c*_j_ = 1 for *j* ∈ L and zero otherwise,

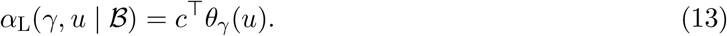

This is an onset contribution fraction. It is not an entry-flux fraction, a read fraction, or ultimate ancestry. Dependence on *u* is suppressed below only when it is unambiguous.

## 4 Identification conditional on a complete scenario

### 4.1 Time-expanded observation map

Fix one environmental–response–model scenario *γ*. The structural certificate first sets *e*_t_ = 0 or conditions on a fixed error realization that has been subtracted. Let *u* contain only unknown history coordinates. Stacking the remaining observations gives the affine map

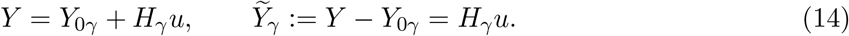

Here *Y*_0γ_ stacks the propagated known initial state and any other fixed history components; it is zero when every history component is included in *u*. Columns for uncertain initial-state coordinates contain 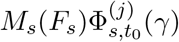. Columns for admissions at time *t < s* contain 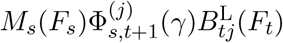 or their imported counterparts and are zero before admission. Known versions of those components belong in *Y*_0γ_. More frequent sampling adds rows. It does not guarantee that local and imported columns become different.

If observation error is unknown but bounded, append its components to *u* and the corresponding identity block to *H*_γ_, with zero coefficients in the source target. The same fiber certificate then applies to the augmented model. Alternatively, optimize the target over the bounded-error fiber. Treating an unknown error as zero without either step is not justified. The onset-channel error in Equation (3) is handled in exactly this way and its threshold constraints remain part of the fiber.

Let 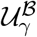 be the convex set of histories allowed by nonnegativity, source support, timing constraints, and the bloom event. Write

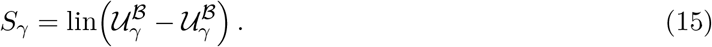

For fixed *γ*, write the target in affine-ratio form:

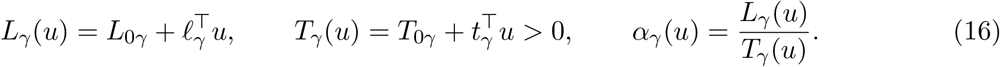

The constants *L*_0γ_ and *T*_0γ_ are the target contributions of the same fixed history components represented in *Y*_0γ_; in particular, fixed initial stock contributes locally to both constants under Equation (7). The vectors *ℓ*_γ_ and *t*_γ_ act on exactly the unknown coordinates used in Equation (14).

For a fixed candidate value *α*, determining (*ℓ*_γ_ −*αt*_γ_)^⊤^*u* from the linear part of the observation record is a finite-horizon functional-observability question [17, 22, 36, 47, 53, 61]. Ordinary full-state or full-input observability is stronger than required for this one target. Here the functional is embedded in a ratio, the admissible directions are restricted by the bloom event and feasible histories, and the complete scenario may itself be uncertain. In the classical structural sense, the certificate concerns ideal noise-free uniqueness under the declared model and experiment, not practical precision and not identification of every biological parameter [8, 46, 68]. The data fiber is the set of source histories that produce the same stacked observations. A null direction is an allowed history change *h* for which *H*_γ_*h* = 0.

#### Theorem 2

(Interior fixed-scenario target certificate). *Suppose* 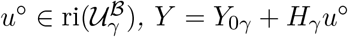, *and α*^°^ = *α*_γ_(*u*^°^). *The target is constant on the observed data fiber*

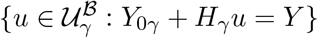

*if and only if*

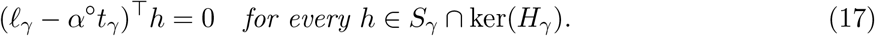

#### Plain-language interpretation

A fixed complete scenario does not make source attribution automatic. Any allowed change in source history that leaves all observations unchanged must also leave the local fraction unchanged. One counterexample direction proves failure.

*Proof*. Relative interior permits sufficiently small perturbations *u*^°^± *εh* for every *h* ∈ *S*_γ_. If *H*_γ_*h* = 0, both perturbations give the same observations. Direct calculation gives

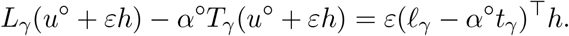

This proves necessity. Conversely, any two points in the fiber differ by an *h* ∈ *S*_γ_ ∩ ker(*H*_γ_). Equation (17) then gives the same ratio because the denominator is positive.

The relative-interior assumption is essential to this clean two-sided null-direction certificate. At a boundary point, nonnegativity or another active inequality may remove one side of a direction. Equation (17) is therefore not a universal pointwise test; the exact general test is equality between the minimum and maximum target over the observed data fiber.

##### Corollary 3

(Non-identification under a shared channel). *Suppose the time-expanded local and imported input blocks are equal. Let h* = (*δ*^⊤^,− *δ*^⊤^)^⊤^ *be a two-sided feasible transfer between local and imported histories. If that transfer changes the local numerator but not the total, then the local fraction is not identified*.

#### Plain-language interpretation

When the instrument sees only the sum of local release and import, mass can be moved between those labels without changing the record. Dense sampling of the same summed channel cannot reveal the split.

*Proof*. The shared channel gives *H*_γ_*h* = 0. The total has zero change, while the local numerator has nonzero change. Theorem 2 applies.

This corollary does not cover source measurements, spatial observations, or input operators that give local and imported carriers distinct observed directions. Those measurements can remove the null direction.

### 4.2 Environmental, response, and model uncertainty

For each *γ* ∈ Γ(*f*), define the feasible data fiber

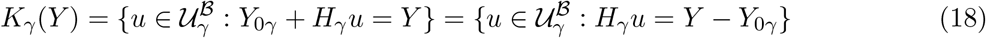

and its scenario-conditional identified set

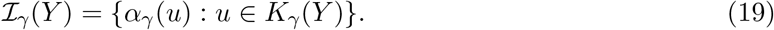

Empty fibers are discarded because they cannot explain the observations. Define the nonempty compatible scenario set

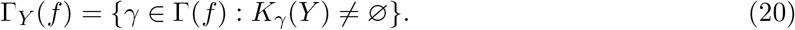

#### Theorem 4

(Identification across compatible scenarios). *Assume* Γ_Y_ (*f*)≠∅, *each K*_γ_(*Y*) *for γ* ∈ Γ_Y_ (*f*) *is compact and convex, and every target denominator is positive. Then each ℐ*_γ_(*Y*) *is a closed interval. The joint identified set is*

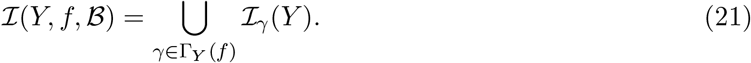

*The local fraction is point identified if and only if every* I_γ_(*Y*), *γ* ∈ Γ_Y_ (*f*), *is the same singleton. For an arbitrary, possibly infinite scenario family, the union in Equation* (21) *need not be an interval or a closed set*.

*For a finite environmental–response–model ensemble, write* 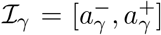. *The sharp lower and upper bounds are*

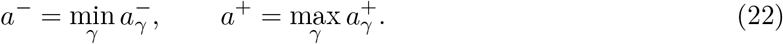

*Let* 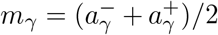 *and* 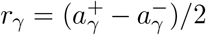. *If γ*_+_ *maximizes m*_γ_ + *r*_γ_ *and γ*_−_ *minimizes m*_γ_ − *r*_γ_, *then the joint bound width is exactly*

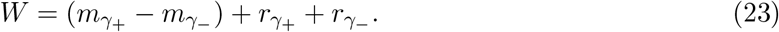

*If R*_m_ = max_γ_ *m*_γ_ − min_γ_ *m*_γ_ *and* 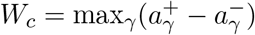, *then*

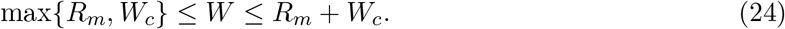

#### Plain-language interpretation

Source-history uncertainty is measured within one complete scenario. Environmental-path, biological-response, and model-form uncertainty move or widen the collection of conditional intervals in distinguishable ways. A point claim is valid only when all compatible scenarios agree on that same point.

*Proof*. A scalar linear-fractional map with positive denominator sends a connected convex set to an interval. Compactness closes the interval and attains its endpoints. Equation (21) follows from allowing every compatible scenario. A union is one point exactly when every nonempty member is that point. For a finite ensemble, the smallest lower endpoint and largest upper endpoint are attainable, proving (22). Writing each endpoint as *m*_γ_ ± *r*_γ_ gives (23). The joint hull contains every conditional interval and every midpoint, giving the lower bound. The midpoint difference is at most *R*_m_, and the two radii sum to at most *W*_c_, giving the upper bound.

#### Exact computation on a polyhedral history fiber

The endpoint existence result above has a direct computational form. Suppose the fixed-scenario history restrictions are polyhedral,

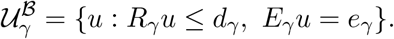

For each nonempty *K*_γ_(*Y*), its dynamic-data endpoints are

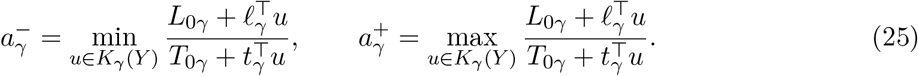

Define the Charnes–Cooper variables *v* = *τu* and 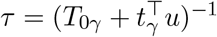 and the transformed polytope

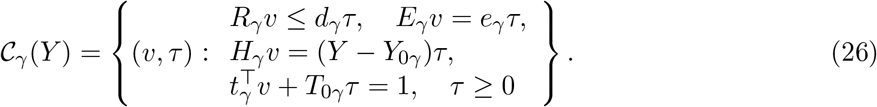

Then the exact endpoints are the two linear programs

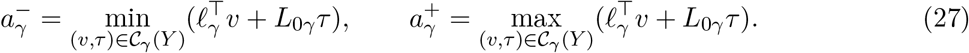

This is the standard linear-fractional transformation [11]. Compactness of *K*_γ_(*Y*) makes its recession cone trivial. A transformed feasible point with *τ* = 0 would therefore force *v* = 0, contradicting the denominator normalization, so *τ >* 0 and *u* = *v/τ* is well defined. The transformation is thus a bijection and the programs are sharp for the full declared dynamic-data fiber. For a finite scenario ensemble, solve these two programs for every compatible scenario and then apply Equation (22). A nonpolyhedral fiber requires the corresponding transformed convex or conic program rather than an LP.

The union in (21) can have gaps and, for an infinite family, can fail to be closed or attain its infimum and supremum. Equations (22)–(24) are finite-ensemble results and give sharp endpoint bounds, not a claim that every intermediate value is feasible. Other scenario families require explicit topological and optimization assumptions. The full union should be reported when its disconnected shape matters.

If Γ_Y_ (*f*) is empty, the declared forcing and source model cannot explain the data. That is model rejection, not point identification.

### 4.3 A forcing–response ambiguity witness

Consider one day between source entry and onset. The local input contains genotype A and the imported input contains genotype B. Each input is 100 carrier equivalents per liter, so the entry mixture is 50:50. The assay observes only total taxon abundance. Under joint scenario *γ*_A_, the genotype A and B gains are (2, 1). Under *γ*_B_, they are (1, 2). Both scenarios produce 300 carrier equivalents per liter. Both cross a predeclared onset threshold of 250 carrier equivalents per liter.

Yet

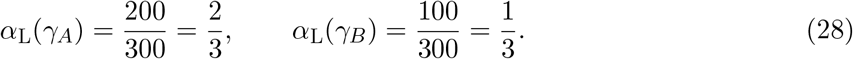

The source mixture and taxon total are identical. Only the genotype-specific response is reversed. Thus taxon-only observations cannot identify the onset fraction when both gain patterns remain compatible. A carrier-resolved assay can separate the scenarios only if the response signatures remain distinct after selection and observation.

Either the environmental path, response parameters, or both can generate the gain reversal, so the witness establishes joint forcing–response scenario ambiguity. It separates source availability from environmental permissibility: identical inputs may remain below onset under nonpermissive forcing, while compatible source histories may differ under a shared permissive path.

## 5 Onset-signature reductions of the dynamic model

**Model hierarchy**. The primary object is the full dynamic-data fiber *K*_γ_(*Y*) over source histories. Each feasible history induces normalized onset signatures *Q*_γ_(*u*). A fixed *Q*_γ_ is justified only when every relevant 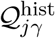 is a singleton after any required class refinement. Otherwise the reduced model must retain 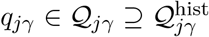 and declare any omitted cross-history or signature–share coupling.

Thus the hierarchy is

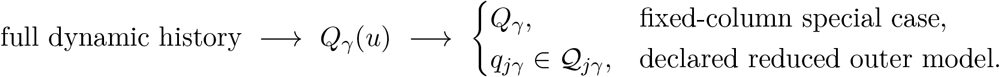

Neither fixing *γ* nor observing onset alone collapses the first model into the fixed-column case.

### 5.1 From labeled states to a source design

After subtracting a fixed onset-observation error realization, suppose the positive total *T*_γ_(*u*) is measured or calibrated on the same carrier scale as the onset assay. For a feasible history *u* with *ϑ*_jγ_(*u*) *>* 0, define the observed response per unit onset contribution:

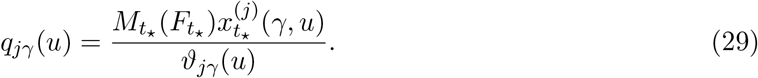

Fixing *γ* fixes the propagation and observation operators. It does not, by itself, fix this normalized column. Different admission times, entry states, or transport routes can give different internal-state compositions at onset.

The exact history-specific scaled identity is

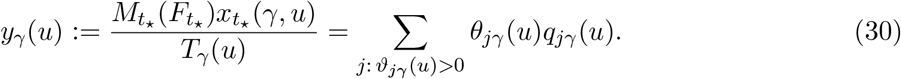

If the onset total is not measured or calibrated, it must remain a nuisance scale and this normalized static design is unavailable as written. A zero-share class may be assigned any admissible column because its product with *θ*_jγ_(*u*) = 0 vanishes; this preserves the fixed dimension *J*. A class that can never contribute should instead be removed or constrained to zero share. After those assignments, set *Q*_γ_(*u*) = [*q*_1γ_(*u*) *q*_Jγ_(*u*)]; then (30) is equivalently *y*_γ_(*u*) = *Q*_γ_(*u*)*θ*_γ_(*u*). Define the history-induced set

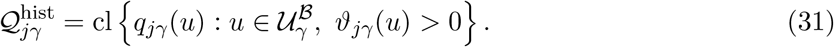

A fixed column is justified only when this set is a singleton, or when class *j* is refined by admission time, entry state, and any other required feature until the normalized signature is fixed. Otherwise Section 6.3 is the default: the signature remains an optimization variable in a set containing 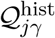.

In the singleton or refined-class case, assign the fixed column also to any zero-share instance, suppress the dependence on *u*, write *Q*_γ_ = [*q*_1γ_ · · · *q*_Jγ_], and obtain, for a realized scaled observation,

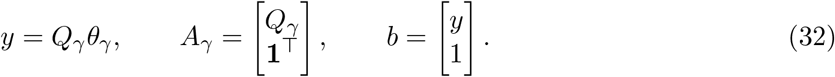

This is the standard convex mixing model *y* = *Qθ, θ* ≥ 0, **1**^⊤^*θ* = 1, encountered in end-member mixing analysis [12, 32], hyperspectral unmixing [10, 15, 23, 45, 48], nonnegative matrix factorization [19, 43], and chemical source apportionment [55]. The mixing law is not new here. The added objects are the forcing-dependent history that induces *Q*_γ_(*u*), recovery of the target *c*^⊤^*θ* rather than necessarily every source share, and identified-set bounds when the source signatures or forcing are uncertain. The history-specific contribution *ϑ*_jγ_(*u*) already contains transport, survival, and growth. Those gains must not be multiplied into *q*_jγ_(*u*) a second time.

Rows may represent calibrated variants, multilocus haplotypes, genome-wide contrasts, or independently measured transport channels. Repeating the same collapsed observation does not add an independent row.

### 5.2 Observation fibers and the quotient

This subsection assumes the singleton or refined-class fixed design above. For a fixed exact design, two source vectors are observationally equivalent when their difference lies in ker(*A*_γ_). The kernel is the set of source changes that the assay cannot see. The quotient ℝ^J^ */* ker(*A*_γ_) retains only distinguishable linear coordinates. The row space is the span of the measured contrast rows. A target lies in that row space when it is a weighted combination of those measured contrasts.

Duplicate source columns are one simple case. More general combinations can also cancel. For example, the columns *q*_1_ = (1, 0)^⊤^, *q*_2_ = (0, 1)^⊤^, and *q*_3_ = (1*/*2, 1*/*2)^⊤^ have no equal pair, but source 3 is observationally equivalent to a mixture of sources 1 and 2.

#### Theorem 5

(Target-specific row-space certificate). *Suppose the feasible set*

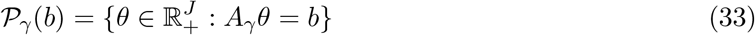

*contains a strictly positive point. The aggregate local fraction c*^⊤^*θ is constant on* P_γ_(*b*) *if and only if*

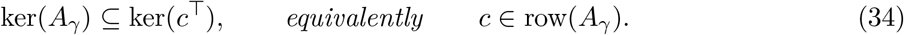

*The complete vector of source–carrier class contributions is identified on all such fibers if and only if* rank(*A*_γ_) = *J*.

**Plain-language interpretation**. The local total can be known even when individual local subclasses are not. Every invisible exchange must stay within one side of the local-import target. Recovering every named source–carrier class requires the stronger full-rank condition.

*Proof*. Let *θ*^°^ *>* 0 be feasible. Every *h* ∈ ker(*A*_γ_) gives feasible points *θ* ± *εh* for small *ε*. The target is constant only if *c*^⊤^*h* = 0 for every such direction. The converse follows because any two feasible points differ by a kernel vector. Orthogonality to the kernel is equivalent to row-space membership. Full source recovery is equivalent to a zero kernel.

The strictly positive feasible point makes this an interior-fiber row-space criterion. On a boundary fiber, sparsity or other active constraints can remove otherwise invisible exchanges, so *c* ∈ */* row(*A*_γ_) does not by itself prove pointwise non-identification. The general boundary test is equality of the minimum and maximum of *c*^⊤^*θ* over the actual feasible fiber.

The row space is the set of target contrasts that can be assembled from the measured contrasts. A source-level contrast need not be scientifically useful. It is useful here only when it spans the local indicator *c*.

### 5.3 Absolute and compositional channels

For calibrated absolute channels, the unit-sum row in *A*_γ_ can provide new information. For a complete compositional profile, 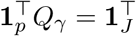. The profile already contains the unit-sum relation. It must not be counted twice.

Choose a contrast matrix *C* ∈ ℝ ^(p−1)×p^ with ker(*C*) = span {**1**_p_}. An independent compositional design is

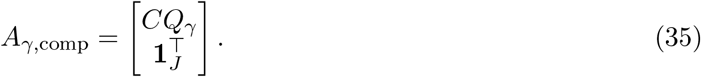

For *J* named source–carrier classes, full recovery needs at least *J* −1 independent composition contrasts plus normalization. The local aggregate may need fewer when its indicator is already in the row space.

An ordinary read-frequency profile weights sources by detection yield. It represents carrier shares only when source-specific yields are equal, calibrated, or included as nuisance transformations.

## 6 Partial identification in onset-signature reductions

### 6.1 Fixed scenario and a fixed source panel

This subsection also assumes history-invariant or refined-class signature columns. If normalized signatures vary across admissible histories, use the uncertain-signature formulation in the next subsection.

Let 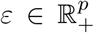 be a deterministic observation tolerance. Let *θ* and 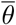 be independently justified source-share bounds. Define

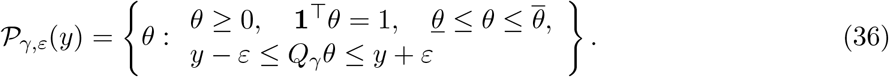

The conditional bounds are

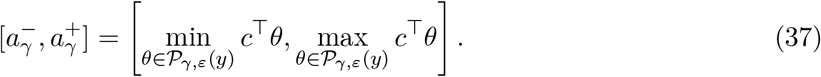

#### Proposition 6

(Sharp fixed-panel bounds). *If P* _γ,ε_(*y*) *is nonempty, the endpoints in* (37) *are attained. Every value between them is produced by at least one feasible source vector*.

#### Plain-language interpretation

The interval contains exactly the local fractions allowed by the fixed complete scenario, fixed source panel, and declared tolerances. The optimization does not add information that was not measured or assumed.

*Proof*. The feasible set is a compact convex polytope. A linear objective attains its minimum and maximum. Its scalar image is the complete interval between those extrema.

These bounds are identified-set bounds, not confidence intervals. Statistical coverage must come from a calibrated joint uncertainty region [34, 38].

### 6.2 From statistical calibration to deterministic identified sets

The tolerance *ε*, environmental-path set Γ_F_ (*f*), response sets Γ_β_(*F*), source caps, onset-channel error set, and signature polytopes are inputs to the deterministic identification analysis; they are not automatically confidence statements. With noisy or estimated inputs, a statistical layer first constructs a calibrated joint uncertainty region ℛ_1−δ_ for the observation record and nuisance objects. The deterministic layer then propagates that region:

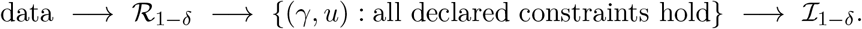

Joint calibration matters because sequencing channels, compositional signatures, environmental trajectories, and fitted response parameters are commonly dependent. Independent coordinatewise tolerances can admit impossible combinations or make bounds unnecessarily conservative. Repeated sampling and replicate assays can tighten the joint region and support model checking even when they do not remove a structural null direction. Formal finite-sample inference is a separate layer; under the maintained model, joint coverage propagates to an outer source-fraction set by set inclusion.

### 6.3 Uncertain source signatures

Unless the fixed-column condition above has been established, this is the default signature reduction. For source–carrier class *j* under forcing scenario *γ*, choose a justified nonempty compact polytope containing its attainable history-induced signatures,

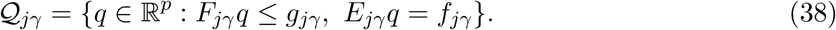

If no bounded envelope containing 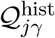 is scientifically justified, this program does not apply. The set may encode admission-time, entry-state, transport-route, assay variation, and within-signature channel dependence. For a compositional signature, impose **1**^⊤^*q* = 1; absolute channels need not sum to one. Complete-scenario uncertainty remains indexed by *γ* rather than being hidden inside one undifferentiated error bar.

Introduce *z*_j_ = *θ*_j_*q*_j_, the observed channel contribution of source *j*. This substitution is a perspective lift. It scales every constraint defining a source signature by that source’s contribution. The scaling removes a bilinear product from the optimization variables.

#### Theorem 7

(Sharp Cartesian-product polyhedral-signature program). *For fixed γ, assume that the maintained admissible joint signature set is exactly the Cartesian product* Q_1γ_ × · · · × Q_Jγ_. *Impose*

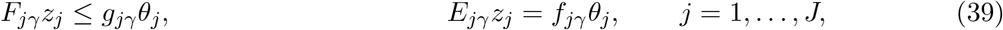

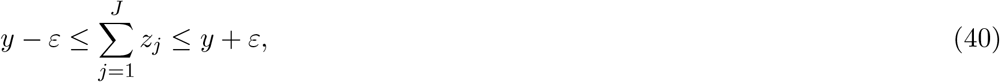

*together with θ* ≥ 0, **1**^⊤^*θ* = 1, *and justified source bounds. Minimizing and maximizing c*^⊤^*θ gives the sharp conditional endpoints over* Q_1γ_ × · · · × Q_Jγ_.

#### Plain-language interpretation

The new variable *z*_j_ records how much of each observed channel came from source *j*. The scaled constraints keep each source signature inside its own uncertainty set. The program searches all allowed signatures without breaking their within-source relationships.

*Proof*. Any feasible (*q*_j_, *θ*_j_) gives *z*_j_ = *θ*_j_*q*_j_ and satisfies (39). If *θ*_j_ *>* 0, the reverse construction is *q*_j_ = *z*_j_*/θ*_j_. If *θ*_j_ = 0, boundedness makes the recession cone of *Q* _jγ_ trivial, so the constraints force *z*_j_ = 0. The lifted program therefore has the same projection onto *θ* as the declared bilinear Cartesian-product signature model. Its target extrema are attained.

Independent coordinate bounds are the rectangular special case. They should not replace known covariance, compositional closure, or trajectory dependence within a signature. Cross-class dependence and signature–share dependence cannot be represented by separate marginal Q_jγ_ sets; they must enter through joint lifted constraints. The term “sharp” is always relative to the maintained uncertainty set. The program is sharp for its declared Cartesian-product signature model. It is also sharp for the underlying dynamic-history model only when the admitted signature and source-share combinations are jointly attainable, or when all history-induced coupling constraints are retained in the lifted program. Marginal outer envelopes give valid but potentially conservative bounds.

For a finite complete-scenario ensemble, Theorem 4 applies directly when each scenario-specific optimization retains every dynamic-data constraint in *K*_γ_(*Y*) and every required history-induced coupling. If only the onset datum *y* and the static envelope are used, define

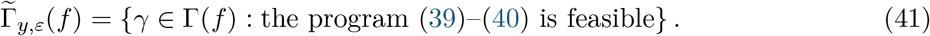

The union of its attainable source fractions is exact for that declared reduced onset-data model. It can be conservative relative to the full dynamic-data set because stacked observations or signature–share couplings were omitted. Marginal outer envelopes can enlarge it further.

### 6.4 Unknown signature and unknown provenance

An omitted source is not zero source mass. If independent evidence establishes that an unknown-signature class is external, its target coefficient is zero while its signature varies over a justified set. If geographic status is unknown, split its contribution into

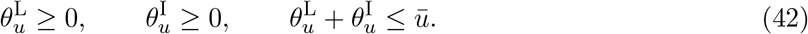

Assign target coefficients one and zero. If the two components share one signature, use one observed vector 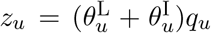. The mass cap 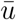 is a sensitivity parameter unless sampling coverage, hydrodynamics, or a controlled experiment supports it. Without that support, the correct bounds may be [0, 1].

## 7 Computational verification

The computations verify equation transcription and expose failure modes in declared synthetic systems; they are mathematical checks rather than field validation. Each example reports its dimensions, carrier scale, source classes, forcing variables, and target interpretation.

Two additional executable controls enforce reporting and schema consistency: a language guard checks generated summaries, and an application-readiness validator checks the declared field-case schema.

An invisible local–import direction is a structural identification failure: regularization may select a point but cannot make it data-identified. An unresolved environmental–response scenario can likewise move the target while the source mixture remains fixed.

## 8 Illustrative field application-readiness cases

Field readiness has two distinct levels. The *full-panel point-claim gate* requires a predeclared control volume, window, carrier scale, and observed or calibrated onset channel; compatible quantitative measurements of every material local compartment and inflow; transport-consistent timing; and jointly calibrated environmental-path, biological-response, model-form, and observation uncertainty. Passing that measurement gate is necessary but not sufficient for a point claim: the full dynamic-history identified set must also be the same singleton for every compatible scenario.

**Table 2.** Synthetic mathematical checks and their scientific purpose.

| Check | Purpose |
| --- | --- |
| Fixed-scenario shared channel | Confirms identical observations and different local fractions along a source null direction. |
| Full-history fractional program | Verifies the affine observation offset and computes both sharp dynamic-data endpoints by the Charnes–Cooper LPs in Equation (27). |
| Joint source–scenario witness | Reproduces the 2/3 versus 1/3 result in Equation (28). |
| Genotype-specific response | Shows that one entry mixture can produce different onset genotype composition under different synthetic environmental paths while the response map and model form remain fixed. |
| Scenario ensemble | Checks conditional intervals, their union endpoints, and the width bound in Equation (24). |
| Taxon versus carrier panels | Returns $[0, 1]$ for an uninformative taxon channel and a narrower set for a separating carrier panel. |
| Unknown provenance | Retains the local-import allocation of an unclassified source rather than assigning it a convenient location. |

**Figure 1:**
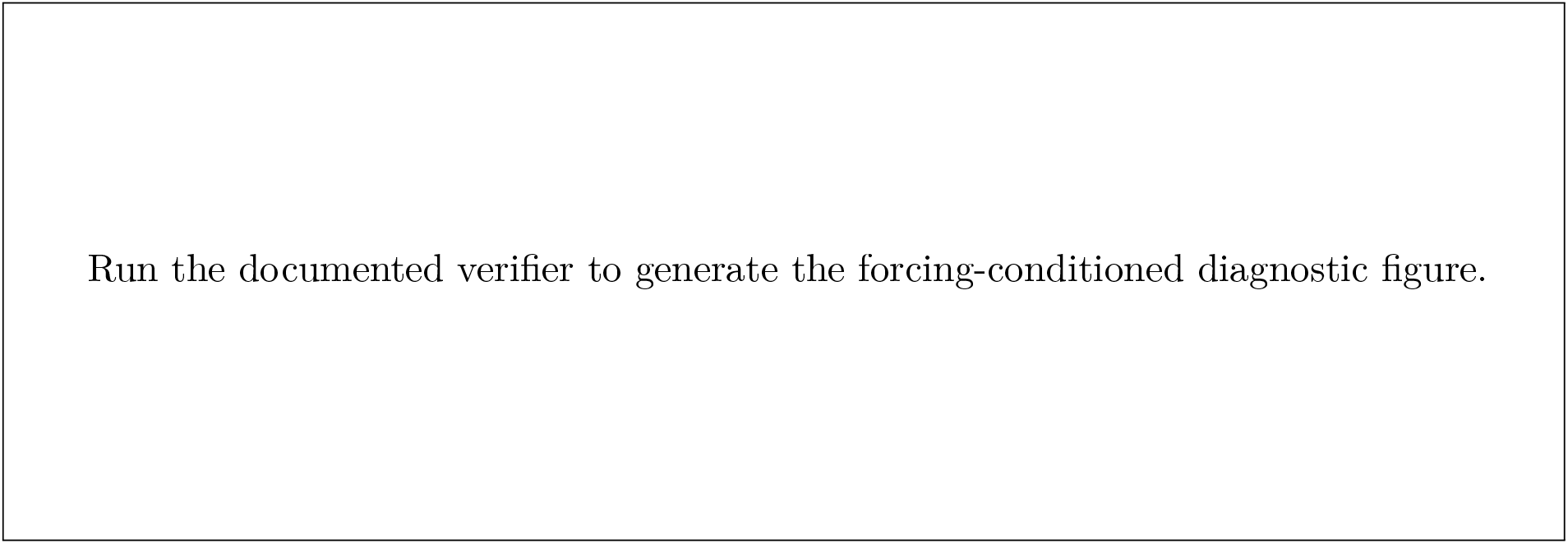
SYNTHETIC DIAGNOSTICS—NO FIELD DATA OR ESTIMATES. (A) As synthetic source signatures alias, the sharp local-fraction interval widens. (B) For one fixed observed onset composition and fixed response parameters/model form, compatible dimensionless environmental paths imply different entry and onset local fractions. (C) With the same 50:50 entry mixture and fixed response map, two complete synthetic scenarios yield different onset genotype compositions. (D) Omitting a synthetic unknown source yields a biased point, whereas a declared mass cap produces an interval; unknown provenance widens it further. The dashed line in D is the synthetic generating value. No panel is calibrated to a biological exposure or field system.

The *partial-identification usability gate* retains the same operational target and onset requirements but permits a missing source, signature, environmental-path or biological-response term, or operator only when each omission has an independently supported joint mass cap, envelope, or operator bound. The resulting sharp set must be nonempty and meet a predeclared informativeness criterion, at minimum being narrower than [0, 1]. Unconstrained missing provenance can yield [0, 1]; in that case neither a preferred fraction nor an informative field interval is reported.

Table 3 applies these gates to an illustrative, nonrepresentative maximum-contrast set of four public-record cases. Nauset Marsh supplies a semi-enclosed geometry, mapped cyst beds, and a dominant inlet; western Lake Erie combines sediment sources, multiple tributary inputs, and extensive monitoring; the Gulf of Maine combines regional seedbeds, population structure, and mature coupled models; LTER-MC supplies a long plankton and genetic record but exposes a source-definition failure. Search endpoints, query families, inclusion and exclusion rules, the cutoff date, and the evidence ledger are documented in the search protocol, 44-record extraction ledger (27 candidate-system and 17 contextual records), case audit, and machine-readable gate schema listed in `SUBMISSION_MANIFEST.md`. Conclusions below are limited to these four cases and the public records examined.

**Table 3.** Illustrative application-readiness cases. No case supports either a full-panel point claim or a calibrated informative field interval.

| System | Useful public evidence | Readiness result |
| --- | --- | --- |
| LTER-MC *P. multistriata* | Foundational long-term plankton record, species life-cycle record, focal genetic time series, hydrodynamic context, and broader genomes [13, 16, 49, 59, 62]. | **Definition stage.** The fixed surface record does not make a predeclared local source class or external boundary operationally observable. The point gate therefore fails before measurement completeness is assessed, and no partial interval is admissible for an undefined target. |
| Nauset Marsh *Alexandrium catenella* | Hydrodynamic, temperature, and residence-time model plus bloom population genetics [58, 60]. | **Full panel: fail; partial interval: not established.** The records do not provide one concurrent common-scale panel for every source, scenario component, and onset population, nor an independent cap or envelope for every omission. |
| Western Lake Erie *Microcystis* | Sediment seed viability and changing *mcy* genotype composition [40, 72]. | **Full panel: fail; partial interval: not established.** External inflows and their transport timing are not closed on a common carrier scale, and no independent omitted-mass/signature envelope supports a nonvacuous interval. |
| Gulf of Maine *Alexandrium catenella* | Cyst maps, regional bloom population structure, and coupled physical-biological source-pathway models [3, 4, 20, 29, 44, 50, 51, 65]. | **Full panel: fail; partial interval: not established.** The reports do not provide a common-assay seedbed-boundary-onset panel or an independent joint envelope for missing sources and scenarios. Mechanistic pathway support is complementary evidence, not a measured carrier fraction. |

Nomenclature follows the molecular grouping of John et al. [37] and the subsequent nomenclatural ruling of Prud’homme van Reine [57]: Group I organisms historically reported as *Alexandrium fundyense* in the North American studies below are referred to here as *A. catenella*. Original article titles in the bibliography retain the names under which they were published.

The Gulf of Naples case remains at the definition stage: under the available fixed-station formulation, the required local class is not observed as an operational source class. Declaring a deeper compartment local after analysis would not repair that design gap.

Across the four illustrative cases, neither the full-panel gate nor the informative-partial-interval gate is cleared. The resulting readiness decisions state what a future design must measure or independently bound; no numerical field fraction is reported.

1. Predeclare the control volume, source window, carrier scale, and onset rule.
2. Observe or calibrate the onset channel and measure the environmental variables supported by the target system’s primary biology, with uncertainty and units.
3. Sample every operational local compartment and material inflow face, or justify an independent joint cap on its omitted contribution.
4. Link source timing to transport or residence-time information.
5. Measure carrier or genotype signatures at sources and onset.
6. Preserve absolute abundance or calibrate source-specific detection.
7. Separate environmental-path, biological-response, model-form, and unknown-source sensitivity.
8. Hold out later composition and timing for model checking.

Source classes must be operationally defined before the theorem can separate them. A defined but unsampled source can enter a partial analysis through an independently supported bound; an undefined class must first be operationalized.

## 9 Relation to previous work

### 9.1 Functional observation, unknown inputs, and parameter identification

Estimating a linear combination of states is the functional-observer problem introduced by Luenberger [47] and developed through existence, eigenspace, rank, and target-sensor conditions [17, 22, 36, 53, 61]. The geometric unknown-input literature asks when unknown disturbances can be made invisible to an observer or reconstructed under strong-observability and detectability conditions [6, 7, 18, 28, 33, 39]. The present fixed-scenario certificate belongs to that lineage. It targets a ratio of two affine functionals on an event-restricted feasible history set, rather than proposing a new full-state or full-input observer.

Structural parameter identifiability asks whether distinct parameter values can produce the same ideal experiment [8, 46, 68]. That is different from asking whether a declared source functional is invariant when parameters or environmental–response scenarios range over an admissible set. The framework can use structurally identified response models, but it does not claim to identify every biological parameter.

### 9.2 Static mixture and partial-identification methods

Microbial source trackers estimate sink mixtures when representative source profiles and a common mixture scale are available [42, 64]. End-member mixing, hyperspectral unmixing, nonnegative factorization, chemical source apportionment, and stable-isotope mixing share the broad nonnegative-mixture geometry discussed after Equation (32). They establish both the long history of the problem and the importance of end-member validity, conservative mixing, separating geometry, and uniqueness. Ecological transformation between source and sink can invalidate a static interpretation [69], and ecological mixing analyses have long reported underdetermination when sources outnumber or overlap the measured contrasts [54, 56].

The row-space condition used here is classical estimability [63]. The distinction is target-specific: the local aggregate may be identified even when all named source shares are not. The perspective lift is also an established optimization device [27]. Robust optimization requires feasibility for every realization in a declared uncertainty set [9]; the identified set here is existential, because each reported value need only be generated by at least one admissible history, signature, and complete scenario. Sampling uncertainty for the resulting population identified set is a separate inferential layer [34, 38, 66].

### 9.3 Comparison with mechanistic HAB and transport models

Benthic-resting-stage studies establish that local cyst or resting-cell reservoirs can seed water-column populations [1–4, 35]. Stage-resolved life-cycle models can explicitly represent germination, encystment, and transitions among vegetative and resting stages [24, 31]. In the Gulf of Maine, coupled physical–biological models have represented cyst germination, coastal-current advection, growth, and event-scale source pathways [29, 44, 50, 51, 65]. Adjoint and coupled models in the Changjiang Estuary likewise identify model-conditional nonlocal source regions whose bloom effects depend on suitable biogeochemical conditions [71]. McGillicuddy et al. [52] shows that downstream physical, biological, and chemical conditions can suppress bloom realization despite an abundant cyst source, anchoring the distinct role of environmental modulation.

Lagrangian analyses and connectivity models use velocity fields, release conditions, and particle behavior to infer pathways, transit times, and connectivity [14, 67]. These are genuine attribution tools for physical delivery. A plausible trajectory or connectivity matrix, however, does not by itself close carrier mass, establish source signatures, or separate delivery from post-entry selection.

Mechanistic trajectory fit and point identification answer different questions: a fitted trajectory is one compatible history, whereas a field point claim requires target invariance across every compatible history and scenario.

**Table 4.**
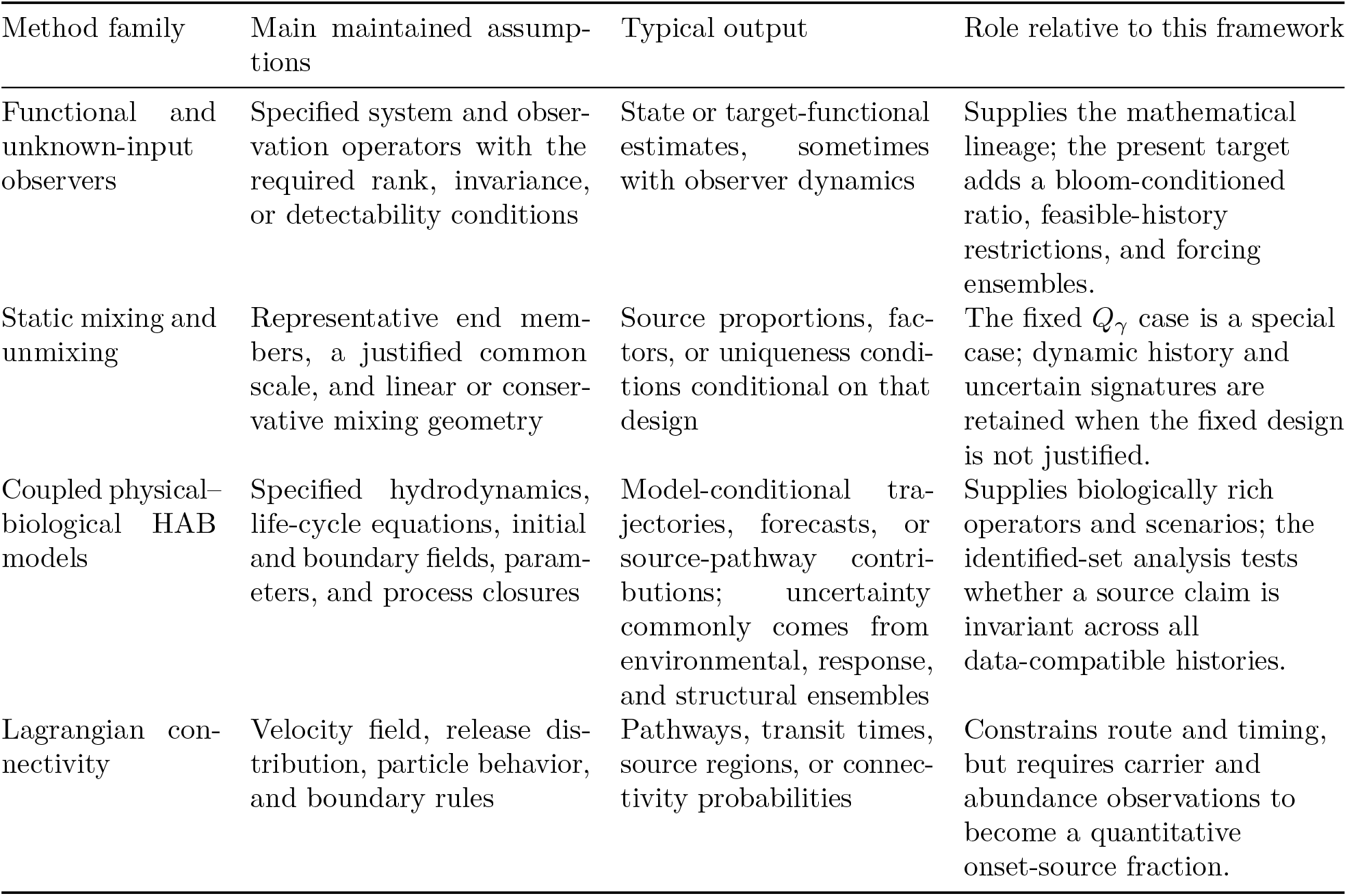
Assumptions, outputs, and complementary roles of relevant method families.

| Method family | Main maintained assumptions | Typical output | Role relative to this framework |
| --- | --- | --- | --- |
| Functional and unknown-input observers | Specified system and observation operators with the required rank, invariance, or detectability conditions | State or target-functional estimates, sometimes with observer dynamics | Supplies the mathematical lineage; the present target adds a bloom-conditioned ratio, feasible-history restrictions, and forcing ensembles. |
| Static mixing and unmixing | Representative end members, a justified common scale, and linear or conservative mixing geometry | Source proportions, factors, or uniqueness conditions conditional on that design | The fixed $Q_\gamma$ case is a special case; dynamic history and uncertain signatures are retained when the fixed design is not justified. |
| Coupled physical–biological HAB models | Specified hydrodynamics, life-cycle equations, initial and boundary fields, parameters, and process closures | Model-conditional trajectories, forecasts, or source-pathway contributions; uncertainty commonly comes from environmental, response, and structural ensembles | Supplies biologically rich operators and scenarios; the identified-set analysis tests whether a source claim is invariant across all data-compatible histories. |
| Lagrangian connectivity | Velocity field, release distribution, particle behavior, and boundary rules | Pathways, transit times, source regions, or connectivity probabilities | Constrains route and timing, but requires carrier and abundance observations to become a quantitative onset-source fraction. |

The approaches are complementary rather than ordered by a universal advantage. A fully specified mechanistic model is richer and can be predictive; its point or ensemble result is conditional on its process, parameter, initial, and boundary assumptions. Partial identification is preferable when the scientific question is what the observations and declared bounds logically determine without collapsing unresolved histories to one fitted solution. It is also a diagnostic wrapper for a mechanistic model: the model supplies *H*_γ_, Γ(*f*), and feasible-history constraints, while the bounds show whether apparent precision comes from the data or from untested source and operator assumptions. The application-specific synthesis combines that diagnostic with a predeclared bloom event and a prospective measurement gate.

## 10 Limitations and scope

1. **Scenario components are system specific**. A forcing vector containing temperature, nutrients, and light is not automatically adequate. Mixing, salinity, grazing, infection, response calibration, or alternative process closures may be material elsewhere and must not be collapsed into one generic uncertainty index.
2. **The onset event is operational**. A threshold crossing defines the analysis time. It is not a universal biological definition of bloom initiation.
3. **Linearity is conditional**. Density dependence, priority effects, cross-source mating, and recombination can break additive source propagation. The source-attribution theorems then do not apply.
4. **Selection changes contributions**. Genotype-specific growth can be represented when labels remain traceable. It cannot be ignored or described as evidence of source origin.
5. **Forcing observations can be endogenous**. Bloom-driven nutrient drawdown or light attenuation requires a coupled state model when material.
6. **Unknown sources remain possible**. A mass cap or signature envelope needs independent support. Otherwise a wide interval is the correct result.
7. **Boundary choice matters**. A deep population may be local under one volume and external under another. Sensitivity to defensible boundaries must be reported.
8. **Reads are not living-carrier counts**. If extracellular DNA or dead cells contribute reads, they require calibration or nuisance channels.

## 11 Conclusion

Environmental forcing and source availability play different roles in a plankton bloom. Forcing governs whether carriers can expand and which carriers are favored. Source histories govern which local and imported carriers are available. A fixed-site record can hide both distinctions.

The local onset fraction is therefore a conditional target. Within a complete scenario, it is identified only when every invisible source-history change leaves that fraction unchanged. Across environmental-path, biological-response, and model-form uncertainty, every compatible scenario must give the same singleton before a point can be reported. Otherwise the correct result is the union of conditional sets or its sharp endpoint bounds. For each polyhedral dynamic-history fiber, those endpoints are obtained exactly with two linear programs.

Genotype-resolved measurements are valuable because post-entry selection can change composition even when source inputs do not change. They are not origin certificates by themselves. A useful point-claim design must measure local sources, external inflow, environmental conditions, biological response, timing, onset, and carriers on one declared window. A partial-identification design may leave components unmeasured only when their joint contribution is independently bounded.

The four illustrative public-record cases did not supply that complete panel or a calibrated nonvacuous partial alternative. Their readiness decisions turn the mathematical result into prospective design requirements: measure every material source or bound its contribution independently, and report a point only when the target is invariant across all compatible histories and scenarios.

## Reproducibility and data availability

The accompanying run folder contains the manuscript source, bibliography, verification code, tests, generated evidence, and literature-search records. The complete field-readiness supplement is enumerated in `SUBMISSION_MANIFEST.md`; it includes the search protocol, narrative audit, 44-record extraction table, machine-readable schema, generated gate decision, source index, and environmental-forcing audit. No new field sequencing data were generated. From the run root, the principal commands are:

~~~
python code\verify_bloom_origin.py ‘
 --output-dir evidence ‘
 --field-schema sources\field\field_schema.json
python -m pytest -q -p no:cacheprovider code
powershell -NoProfile -ExecutionPolicy Bypass ‘
  -File manuscript\source\build_tex.ps1
~~~

The build produces `manuscript/rendered/Conditional_Bloom_Onset_Source_Attribution.pdf`. The verifier is deterministic and records its check and test counts in the run summary. Exact counts should be taken from that generated summary rather than copied into the scientific claim.

## A Notation and interpretation

**Table 5.** Principal objects and their roles.

| Symbol | Meaning |
| --- | --- |
| $B$ | Predeclared three-dimensional control volume. |
| $[t_0, t_\star]$ | Source window ending at the predeclared onset event. |
| $F_t$ | System-specific environmental and ecological forcing state. |
| $\beta_j$ | Response parameters for carrier class $j$. |
| $\mu$ | Declared model-form and operator-family index. |
| $\gamma = (F, \beta, \mu)$ | One complete environmental-path, biological-response, and model-form scenario. |
| $\Gamma(f)$ | Complete scenarios compatible with the environmental observations and declared response and model-form sets. |
| $x_t^{(j)}$ | State contribution carrying source-event label $j$. |
| $a_{tj}, m_{tj}$ | Local release and imported inflow histories. |
| $T_\gamma(u)$ | Positive total onset-carrier contribution. |
| $\vartheta_{j\gamma}(u)$ | Unnormalized onset contribution of source-carrier class $j$ under history $u$. |
| $\theta_{j\gamma}(u)$ | Normalized onset contribution of source-carrier class $j$ under history $u$. |
| $\alpha_L$ | Aggregate local onset contribution, conditional on a complete scenario and the bloom event. |
| $N_t^{\text{obs}}$ | Observed or calibrated onset channel, including its declared joint error set. |
| $Y_{0\gamma}$ | Time-expanded observation offset from known initial state and other fixed history components. |
| $H_\gamma$ | Linear part of the affine time-expanded map from unknown source histories to observations. |
| $Q_\gamma(u)$ | History-specific onset source-signature matrix under scenario $\gamma$; written $Q_\gamma$ only after invariance or class refinement. |
| $\mathcal{Q}_{j\gamma}^{\text{hist}}$ | Closed envelope of history-induced attainable onset signatures for class $j$ under scenario $\gamma$. |
| $\mathcal{I}_\gamma$ | Source identified set conditional on one complete scenario. |

## B Linear-program variables

For a dynamic history vector *u* ∈ ℝ^n^, the full-data Charnes–Cooper program uses *n*+1 core variables (*v, τ*). Equations (26)–(27) homogenize every history, event, and affine-observation constraint, then normalize the positive target denominator. Solving the minimizing and maximizing LPs returns the endpoint histories as *u* = *v/τ*.

Separately, for a reduced onset-signature model with *p* observed channels and *J* sources, the polyhedral perspective program uses *J* + *pJ* core variables:

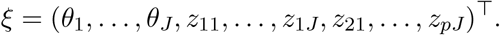

The objective is *c*^⊤^*θ*. Each channel supplies two observation inequalities. Source *j* supplies the scaled inequalities and equalities in (39). A compositional source also satisfies **1**^⊤^*z*_j_ = *θ*_j_.

For a finite complete-scenario ensemble, the appropriate full-history or deliberately reduced program is solved twice per scenario. The scenario-specific minima and maxima feed Equation (22). This keeps source uncertainty within a scenario separate from environmental–response–model uncertainty across scenarios; the reduced signature program is not substituted for the full-history program unless its omitted constraints and any resulting conservatism are declared.

## C Checklist for a future field attribution

1. Is the bloom-onset rule fixed before source analysis and tied to an observed or calibrated channel?
2. Are the material environmental variables measured, estimated, or bounded?
3. Are genotype-specific responses and material model-form alternatives represented or bounded?
4. Are the local and external source classes operationally measurable?
5. Does the source window match transport and residence-time information?
6. Are carrier signatures connected to a common abundance scale?
7. Can source-event labels propagate additively over the window?
8. Are environmental-path, biological-response, model-form, and source uncertainty reported separately?
9. Are unknown sources and unknown geographic status retained?
10. Is the reported quantity clearly separated from bloom causation and genealogical ancestry?

## AI and Programmatic Research Provenance Statement

The originating hypothesis, initial formulation, and initial structure of this work were generated and provided by UCP Technology LLC through its programmatic research process. The source record for the originating research candidate and subsequent development is maintained at: https://github.com/joe-ucp/Luigi2

Artificial intelligence was used as a computational component of that process to generate and develop the originating research candidate, including the initial theoretical formulation, research development, manuscript drafting, literature-supported development, and computational verification.

Artificial intelligence was not used to provide the independent scientific judgment determining whether the work was correct, useful, novel, or suitable for publication.

The human author independently evaluated, revised, verified, interpreted, and approved the final work.

